# Tracing the evolutionary history and global expansion of *Candida auris* using population genomic analyses

**DOI:** 10.1101/2020.01.06.896548

**Authors:** Nancy A. Chow, José F. Muñoz, Lalitha Gade, Elizabeth Berkow, Xiao Li, Rory M. Welsh, Kaitlin Forsberg, Shawn R. Lockhart, Rodney Adam, Alexandre Alanio, Ana Alastruey-Izquierdo, Sahar Althawadi, Ana Belén Araúz, Ronen Ben-Ami, Amrita Bharat, Belinda Calvo, Marie Desnos-Ollivier, Patricia Escandón, Dianne Gardam, Revathi Gunturu, Christopher H. Heath, Oliver Kurzai, Ronny Martin, Anastasia P. Litvintseva, Christina A. Cuomo

## Abstract

*Candida auris* has emerged globally as a multidrug-resistant yeast that can spread via nosocomial transmission. An initial phylogenetic study of isolates from Japan, India, Pakistan, South Africa, and Venezuela revealed four populations (Clades I, II, III, and IV) corresponding to these geographic regions. Since this description, *C. auris* has been reported in over 30 additional countries. To trace this global emergence, we compared the genomes of 304 *C. auris* isolates from 19 countries on six continents. We found that four predominant clades persist across wide geographic locations. We observed phylogeographic mixing in most clades; Clade IV, with isolates mainly from South America, demonstrated the strongest phylogeographic substructure. *C. auris* isolates from two clades with opposite mating types were detected contemporaneously in a single healthcare facility in Kenya. We estimated a Bayesian molecular clock phylogeny and dated the origin of each clade within the last 339 years; outbreak-causing clusters from Clades I, III, and IV originated 34-35 years ago. We observed high rates of antifungal resistance in Clade I, including four isolates resistant to all three major classes of antifungals. Mutations that contribute to resistance varied between the clades, with Y132F in *ERG11* as the most widespread mutation associated with azole resistance and S639P in *FKS1* for echinocandin resistance. Copy number variants in *ERG11* predominantly appeared in Clade III and were associated with fluconazole resistance. These results provide a global context for the phylogeography, population structure, and mechanisms associated with antifungal resistance in *C. auris*.

**Importance:** In less than a decade, *C. auris* has emerged in healthcare settings worldwide; this species is capable of colonizing skin and causing outbreaks of invasive candidiasis. In contrast to other *Candida* species, *C. auris* is unique in its ability to spread via nosocomial transmission and its high rates of drug resistance. As part of the public health response, whole-genome sequencing has played a major role in characterizing transmission dynamics and detecting new *C. auris* introductions. Through a global collaboration, we assessed genome evolution of isolates of *C. auris* from 19 countries. Here, we described estimated timing of the expansion of each *C. auris* clade and of fluconazole resistance, characterized discrete phylogeographic population structure of each clade, and compared genome data to sensitivity measurements to describe how antifungal resistance mechanisms vary across the population. These efforts are critical for a sustained, robust public health response that effectively utilizes molecular epidemiology.

## Introduction

In the last decade, *Candida auris* has emerged in healthcare settings as a multidrug-resistant organism in over 30 countries worldwide (1). Primarily a skin colonizer, this pathogenic yeast can cause bloodstream infections and other infections (2), is often resistant to multiple classes of antifungal drugs (3, 4), and can spread via nosocomial transmission causing outbreaks of invasive infections (5–12).

Initial studies suggested that *C. auris* emerged simultaneously and independently in four global regions, as phylogenetic analyses revealed four major clades of *C. auris* wherein isolates clustered geographically (13). These clades are referred to as the South Asian, East Asian, African, and South American clades or clades I, II, III, and IV, respectively (14). The isolates from these clades are genetically distinct, differing by tens to hundreds of thousands of single nucleotide polymorphisms (SNPs), with nucleotide diversity nearly 17-fold higher between clades compared to within clades (14). All the clades, except Clade II, have been linked to outbreaks of invasive infections; uniquely, Clade II appears to have a propensity for ear infections (15). The need for increased global efforts to understand the population structure of *C. auris* was recently highlighted by the discovery of the first Iranian *C. auris* case that yielded a single isolate representing a fifth major clade (16).

Molecular epidemiologic investigations of *C. auris* outbreaks generally show clusters of highly related isolates, supporting local and ongoing transmission (7, 17, 18). The analysis of outbreaks and individual cases has also revealed genetic complexity, with isolates from different clades detected in Germany (19), United Kingdom (20) and United States (21), suggesting multiple introductions into the countries, followed by local transmission. To date, each of the clades appears to have undergone clonal expansion; while *C. auris* genomes have conserved mating and meiosis genes, only one of the two fungal mating types are present in a given clade. Specifically, *MTL***a** is present in Clades I and IV and the other mating type, *MTL*α, is found in Clades II and III (14). Understanding whether mating and recombination between clades is occurring is critical, especially in those countries, where isolates from different clades and opposing mating types overlap in time and space. This information could help contextualize complex epidemiologic findings or transmission dynamics.

In addition to its transmissibility, *C. auris* is concerning because of its high rates of drug resistance. Three major classes of antifungal drugs are currently approved for systemic use – azoles, polyenes, and echinocandins. Most *C. auris* isolates are resistant to fluconazole (azole) (22). Elevated minimal inhibitory concentrations (MICs) to amphotericin B (polyene) have been reported in several studies and resistance to echinocandins is emerging in some countries (22). Numerous mechanisms of antifungal resistance have been described for *C. auris*. Echinocandin resistance has been linked to a single mutation at S639P/F in *FKS1*, the gene that encodes the echinocandin target 1,3-beta-D-glucan synthase (23). Most isolates display a mutation linked to fluconazole resistance in *C. albicans*; three mutations, Y132F, K143R, and F126L, have been identified in *ERG11*, the gene that encodes the azole target lanosterol 14-α-demethylase. These mutations have been shown to associate by clade where Y132F and K143R are predominately found in Clades I and IV and F126L is exclusively in Clade III (13, 24). Additionally, there have been suggestions that increased copy number of *ERG11* may be a mechanism for fluconazole resistance in *C. auris* (14).

To better understand *C. auris* emergence and population structure, we engaged in a global collaboration involving 19 countries to produce a large dataset of *C. auris* whole-genome sequences from hundreds of cases and associated environmental samples from healthcare surfaces. Our goal was to generate a comprehensive genomic description of a global *C. auris* population to provide a population genetic framework for the molecular epidemiologic investigations.

## Results

### Geographic distribution of *C. auris* major clades

We first performed a phylogenetic analysis to characterize the global distribution of *C. auris* clades. By including isolates from previous studies (13, 18), we observed that all 304 isolates in this sample collection clustered in one of the four major *C. auris* clades (**Figure 1A**). In this collection, 126 (41%) were classified as Clade I, seven (2%) as Clade II, 51 (17%) as Clade III, and 120 (39%) as Clade IV. Globally, Clade I was the most widespread and found in ten countries (Canada, France, Germany, India, Kenya, Pakistan, Saudi Arabia, United Kingdom, United Arab Emirates, and United States); Clade II was found in Canada, Japan, South Korea, and United States; Clade III in Australia, Canada, Kenya, South Africa, Spain, and United States; and Clade IV in Colombia, Israel, Panama, United States, and Venezuela (**Figure 1B**). Multiple clades were found in Canada, Kenya, and United States.

**Figure 1.**
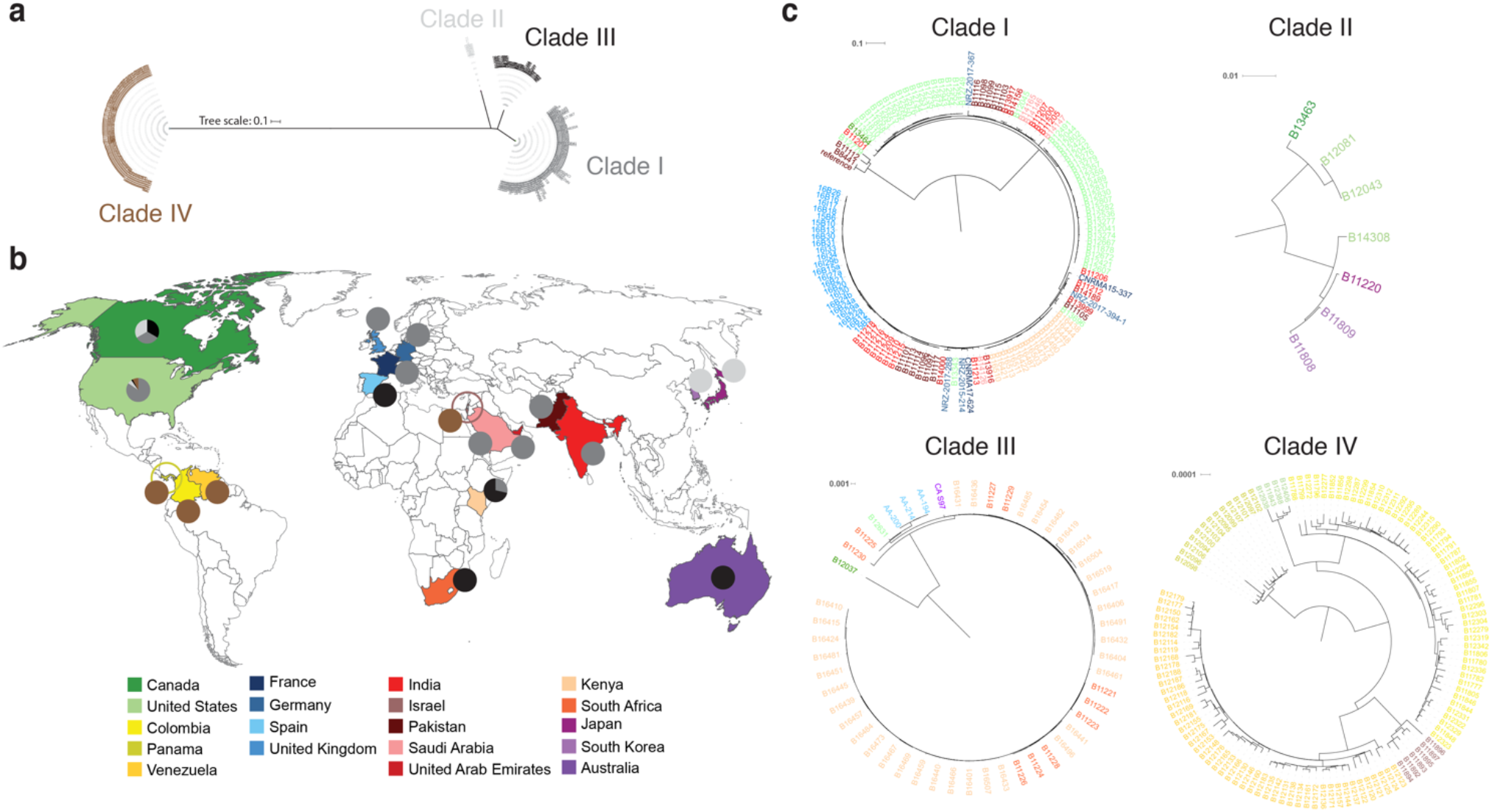
Global distribution of *Candida auris* clades. **(a)** Phylogenetic tree of 304 *C. auris* whole-genome sequences clustering into four major clades. Maximum likelihood phylogeny using 222,619 SNPs based on 1,000 bootstrap replicates. **(b)** Map detailing *C. auris* clade distribution by country (*n* = 19). **(c)** Phylogenetic tree of Clades I to IV; colors represent country.

In contrast to an initial report (13), we observed a weaker phylogeographic substructure as isolates from countries of most global regions appeared interspersed in phylogenies, although there was notable clustering by country within Clade IV (**Figure 1C**). Within Clade I, there were three predominant subclades, each including isolates from India and Pakistan. The smallest subclade included the B8441 reference genome and three other isolates. The other two subclades were more closely related to each other and included groups of highly related isolates from outbreaks in Kenya, United Kingdom, and United States. Additionally, Clade I isolates from countries in Europe (Germany, France) and the Middle East (Saudi Arabia, United Arab Emirates) appeared interspersed in the phylogeny, suggesting multiple introductions of *C. auris* into these countries. Clade II was rarely observed and consisted of seven diverse isolates from Japan, South Korea, United States, or Canada, where six were from cases involving ear infections. Other examples of phylogeographic mixing included isolates from Australia and Spain clustered with Clade III and isolates from Israel clustered with Clade IV, clades originally described as the African and South American clades, respectively.

### Evolutionary rate and molecular dating

To better understand the emergence of this species, we next estimated the divergence times of the four major clades. We utilized collection dates for clinical isolates and associated environmental samples, such as swabs from healthcare facilities, which ranged from 2004 to 2018; most (98%) were collected from 2012 to 2018 (**Figure 2A**). By analyzing Clade III isolates from a single healthcare facility in Kenya experiencing an outbreak of ongoing transmission (25), we confirmed that the divergence level of these isolates are temporally corelated, supporting use of molecular clock analyses, and calculated a mutation rate of 1.8695^e-5^ substitutions per site per year (**Supplementary Figure 1**). This rate was used for a Bayesian approach for molecular dating of a phylogeny for all four clades. Using a strict clock coalescent model, we estimated that the time to most recent common ancestor (TMRCA) for each clade occurred within the last 339 years (**Figure 2B, C**). Clade IV emerged most recently with a TMRCA of 1984 (95% HPD 31.5 - 37.1 years ago), while Clade II was the oldest with a TMRCA of 1679 (95% HPD 321.2 - 356.4 years ago). Lastly, we observed the divergence within the 19^th^ century for the two most closely related clades, Clades I and III, 1878 (95% HPD 132.8 - 149.0 years ago) and 1844 (95% HPD, 164.2 - 187.3 years ago), respectively (**Figure 2B, C**). These dates are impacted by the inclusion of divergent isolates in both clades, which notably do not have *ERG11* resistant mutations and are often drug susceptible, one isolate from Canada in Clade III and two isolates from Pakistan and one from United States in Clade I (**Figures 1C** and **2B**). Excluding these drug susceptible outliers for Clades I and III, TMRCA estimates are for Clade I in 1984 (95% HPD, 17.5 – 21.74 years ago) and for Clade III in 1985 (95% HPD, 16.0 – 21.5 years ago); these more recent estimates are more similar to that estimated for Clade IV. The estimated dates of TMRCA for each individual clade supports the recent expansion during the ongoing outbreak. Together this suggests an older separation of the four clades and the recent diversification of each clade in the years before the detected outbreaks.

**Figure 2.**
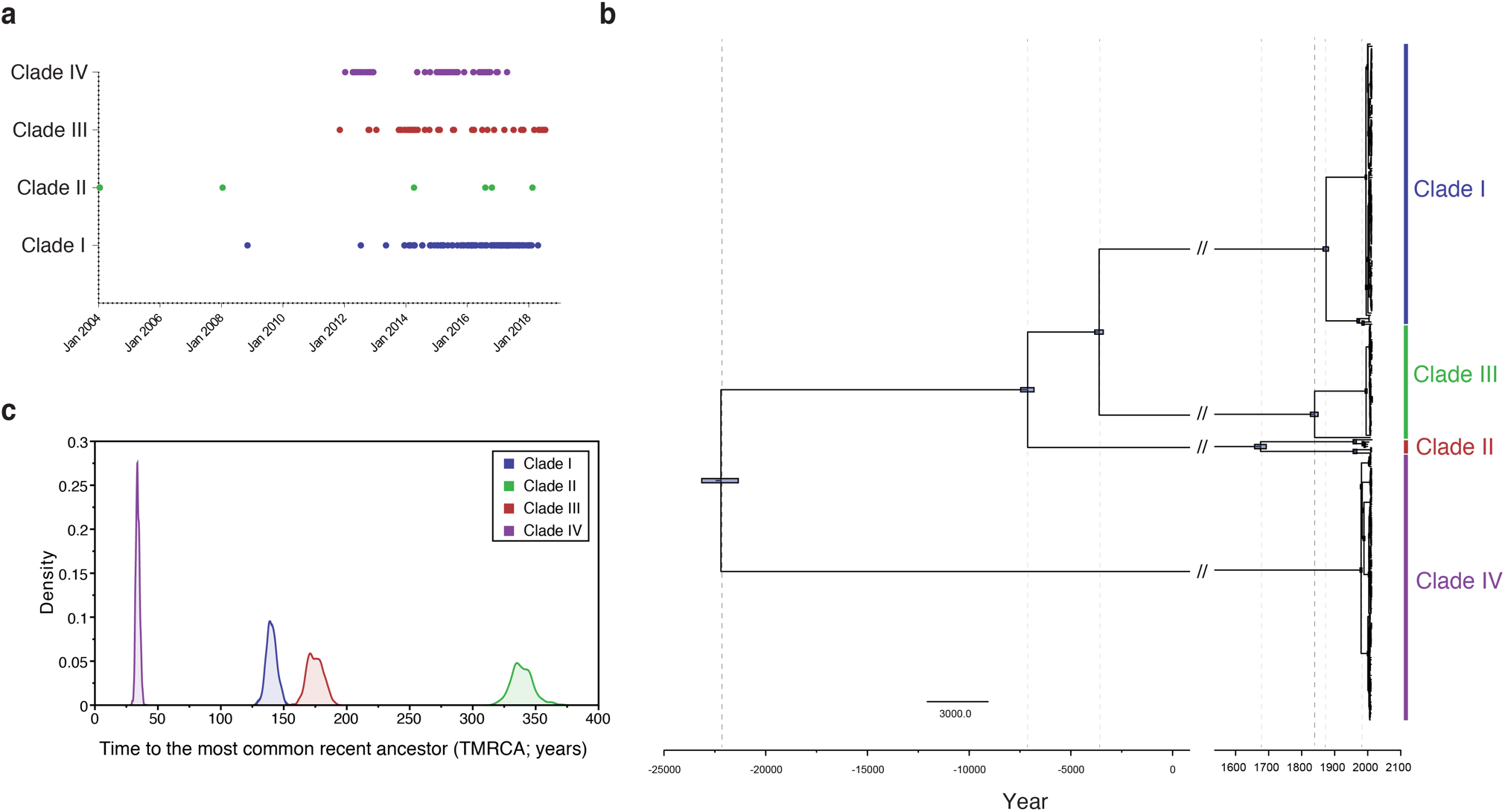
Dating the emergence of *Candida auris*. **(a)** Distribution of collection dates for source specimens by clade. **(b)** Maximum clade-credibility phylogenetic tree of *C. auris* estimated using BEAST (strict clock and coalescent model). Purple bars indicate 95% highest probability density around a node. **(c)** Marginal posterior distributions for the date of the most recent common ancestor (TMRCA) of *Candida auris* Clades I, II, III and IV. The Bayesian coalescent analysis was performed with BEAST.

### *C. auris* population structure

We next examined the global genomic data set for evidence of population substructure and recent admixture. Principal Component Analysis (PCA) identified four well separated populations corresponding to Clades I, II, III, and IV, and the tight clustering of isolates within each clade suggests there is no recent admixture in any isolates (**Figure 3A**). To assess genetic diversity at the clade level, we compared population genetic statistics, including nucleotide diversity (π), Tajima’s D (TD), fixation index (F_ST_) and pairwise nucleotide diversity (D_XY_) (**Figure 3B** to **E**). Overall, Clade I and III showed the lowest genetic diversity (π = 1.51e-5 and π = 1.42e-5, respectively); Clade IV exhibited nearly three times these levels (π = 4.23e-5) and Clade II presented the highest genetic diversity (π = 1.29e-4), nearly nine times higher than Clades I and III (**Figure 3B**). Clade II was also the only clade that exhibited positive TD (td = 1.153; **Figure 3B** and **C**), suggesting balancing selection as expected from the long branches observed in Clade II phylogeny (**Figure 1C**). In Clades I, III, and IV we observed negative TD values consistent with recent population expansions and the shorter phylogenetic branches (**Figure 3B** and **C**, **Figure 1C**); however, Clade IV exhibited a highly variable distribution of TD relative to Clade I and III (**Figure 3C**; **Supplementary Figure 2**), which suggests that these clades have experienced distinct evolutionary processes, such as different degrees of population bottlenecks.

**Figure 3.**
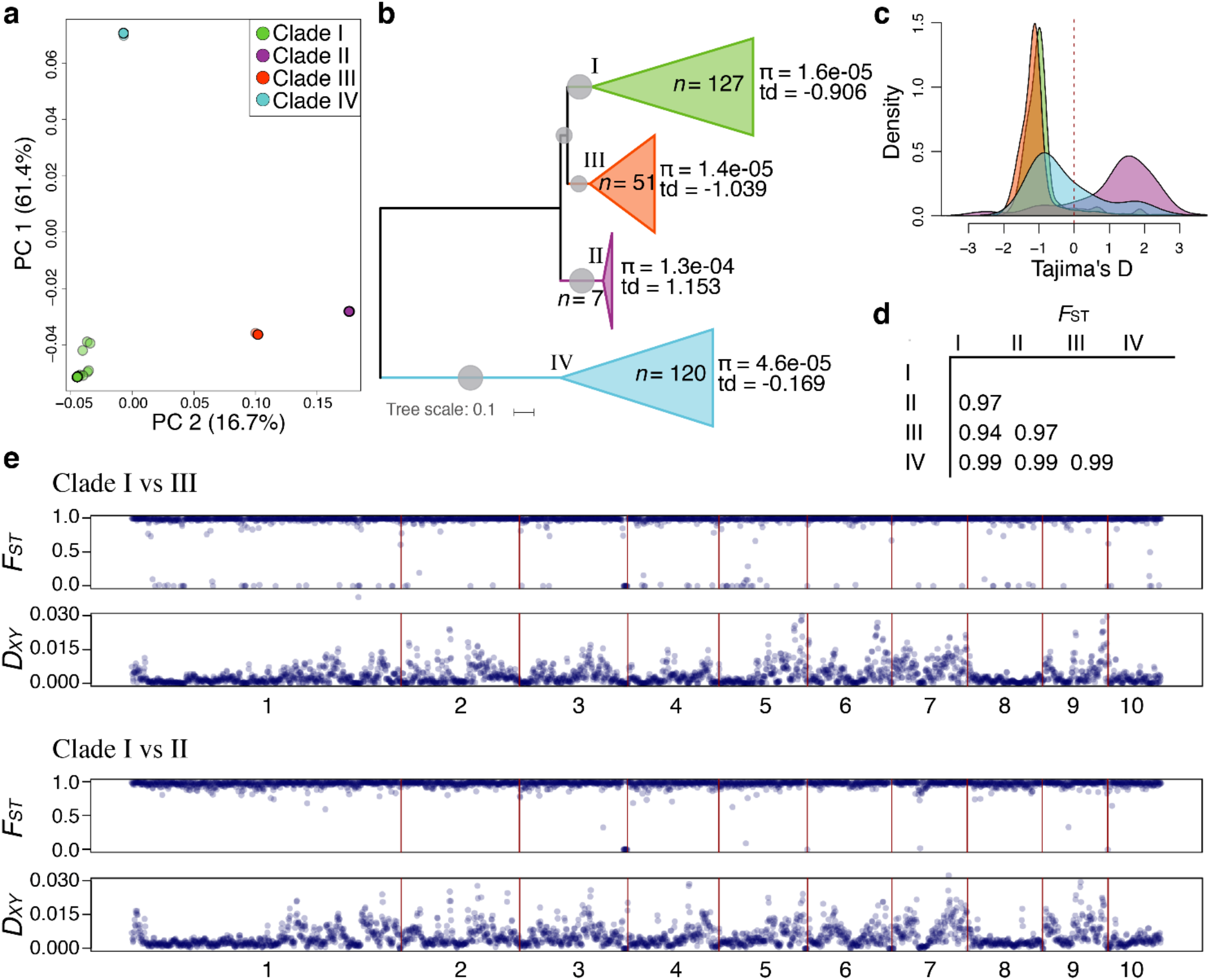
Population structure and genetic differentiation in *Candida auris*. **(a)** PCA analysis and **(b)** phylogenetic tree of 304 *C. auris* depicting genome-wide population genetic metrics of nucleotide diversity (π) and Tajima’s D (TD) for each clade. **(c)** Genome-wide distribution of TD for each clade. **(d)** Average of genome-wide (5-kb windows) variation in fixation index (F_ST_), for pairwise comparisons in each clade as designated in the first vertical and horizontal row. **(e)** Genome-wide (5-kb windows) pairwise FST and pairwise nucleotide diversity (D_XY_) between Clade I and Clade III and Clade I and Clade II are show across the ten largest scaffolds of the B8441 reference genome. All pairwise comparisons of π, TD, F_ST_ and D_XY_ are shown in **Supplementary Figure 2**.

Genome-wide F_ST_ analysis highlighted substantial interspecific divergence and reproductive isolation between *C. auris* clades (average genome-wide F_ST_ > 0.94 in all inter-clade comparisons; **Figure 3D**; **Supplementary Figure 2**). Comparison of the two most closely related clades (Clades I and III) revealed small regions with F_ST_ values close to zero; these regions of identity were distributed across the genome (5.86% of the genome; **Figure 3E**; **Supplementary Figure 2, 3**). Phylogenetic analysis revealed that these regions in isolates from Clades I and III are intermixed in a monophyletic clade (**Supplementary Figure 3**). Comparison of D_XY_ values across the genome highlighted regions of population divergence between *C. auris* clades. Even between these clades with substantial interspecific divergence, we detected large genomic tracts that exhibit either high or low D_XY_ values. For D_XY_, we observed that all chromosomes exhibited a bimodal distribution of regions of both high and low levels of D_XY_; scaffolds 8 and 10, which correspond to chromosomes 1 and 3, respectively (14), show only low D_XY_ values (**Figure 3E**; **Supplementary Figure 2**). D_XY_ is expected to be elevated in regions of limited gene flow, which could have arisen in *C. auris* due to chromosomal rearrangements between the clades (26, 27), whereas D_XY_ is unchanged or decreased in regions under recurrent background selection or selective sweeps.

We observed a single *C. auris* mating type in each clade. Isolates in Clades I and IV had *MTL***a** and those in Clades II and III had *MTL*α (**Supplementary Figure 4**); this confirms prior findings from a smaller data set (14) in this larger global survey. Countries with multiple clades (i.e., Canada, Kenya, and United States) had isolates of opposite mating types; however, there is no evidence of hybridization between clades within these countries or even between isolates of opposite mating types that were observed contemporaneously in a single healthcare facility in Kenya based on the PCA analysis. Together, these findings suggest that the *C. auris* clades have been genetically isolated and that variation across the genome was likely impacted by karyotype variation that prevented equal chromosome mixing.

**Figure 4.**
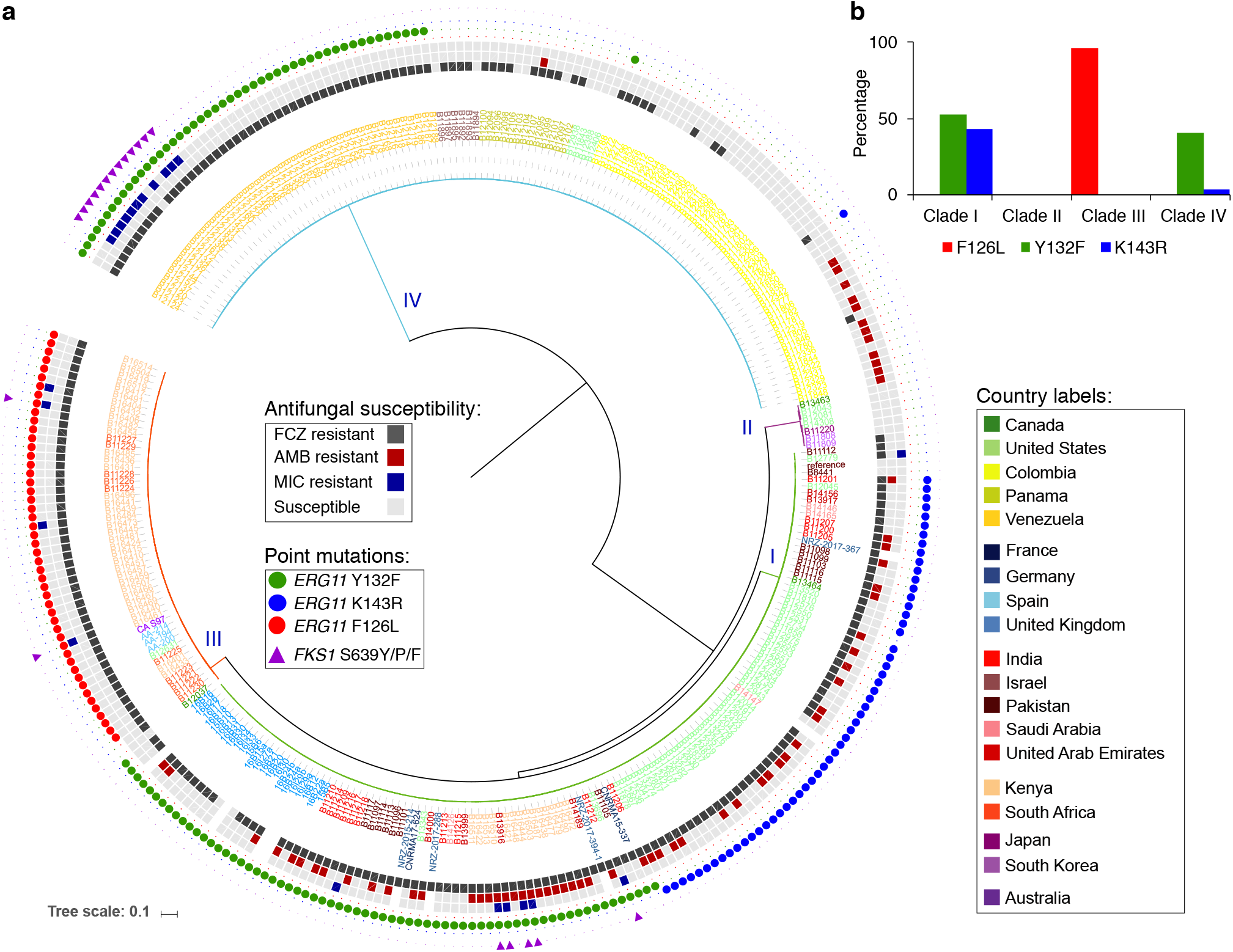
Antifungal susceptibility and point mutations in drug targets in *Candida auris*. **(a)** Phylogenetic tree detailing clade, susceptibility to fluconazole, amphotericin B, micafungin, and point mutations in *ERG11* (Y132F, K143R, F126L) and *FKS1* (S639Y/P/F) associated with resistance. **(b)** Bar plot describing frequency (%, y-axis) of Y132F, K143R, F126L point mutations in *ERG11* for each clade.

### Antifungal drug resistance and mechanisms of resistance

To examine resistance levels, we performed antifungal susceptibility testing (AFST) to fluconazole, amphotericin B, and micafungin – drugs representing each of the major classes. Of the 294 isolates tested, 80% were resistant to fluconazole, 23% to amphotericin B, and 7% to micafungin (**Table 1**, **Figure 4**). Clade II had the greatest percentage (86%) of susceptible isolates, including only one isolate resistant to fluconazole, and Clade I had the greatest percentage of resistant isolates to fluconazole (97%) and amphotericin B (54%). Additionally, Clade I had the highest rates of multidrug-resistance (two antifungal classes; 49%) and was the only clade to have extensive drug-resistance (3%) to all three major classes of antifungals, including isolates from two geographic regions (United Arab Emirates and Kenya) that cluster together (**Table 1**; **Figure 4**). Amphotericin B resistance only appeared in Clades I and IV, and was dispersed across the phylogeny in Clade I and detected in a Clade IV cluster of isolates from Colombia. Clade IV also had the highest percentage (9%) resistant to micafungin, all isolates from Venezuela. Micafungin resistance appeared sporadically in the phylogenies of Clades I and III.

**Table 1.** Frequency of antifungal drug resistance among *Candida auris* isolates by clade.

|  | **Clade I**<br>N=116<sup>a</sup> | **Clade II**<br>N=7 | **Clade III**<br>N=51 | **Clade IV**<br>N=120 | **Total**<br>N=294 |
| --- | --- | --- | --- | --- | --- |
|  | % (n) | % (n) | % (n) | % (n) | % (n) |
| **Susceptible** | 6 (7) | 86 (6) | 2 (1) | 30 (36) | 17 (50) |
| **Fluconazole resistant** | 97 (112) | 14 (1) | 98 (50) | 59 (71) | 80 (234) |
| **Amphotericin B resistant** | 47 (54) | 0 (0) | 0 (0) | 11 (13) | 23 (67) |
| **Micafungin resistant** | 6 (7) | 0 (0) | 8 (4) | 9 (11) | 7 (22) |
| **MDR<sup>b</sup>** | 49 (57) | 0 (0) | 8 (4) | 10 (12) | 24 (72) |
| **XDR<sup>c</sup>** | 3 (4) | 0 (0) | 0 (0) | 0 (0) | 1 (4) |

<sup>a</sup> Complete AFST data for 10 of the 126 (3%) Clade I isolates was missing
<sup>b</sup> MDR indicates multidrug-resistance to two major antifungal classes
<sup>c</sup> XDR indicates extensive drug-resistance to three major antifungal classes

We next determined the genotype of specific drug mutations in the *ERG11* gene that have been associated with azole resistance (Y132F, K143R and F126L). The most widespread mutation was Y132F spanning 11 countries in 53% of isolates from Clade I and 40% of isolates from Clade IV (**Figure 4**). *ERG11* K143R was predominately found in a subclade within Clade I (43%) and one isolate from Clade IV. F126L was found only in Clade III, in nearly all isolates (96%) (**Figure 4**); all isolates with F126L also carried the adjacent mutation V125A. Nearly all of the isolates with these changes in *ERG11* were resistant to fluconazole; 99% of the isolates with Y132F or K143R and 100% of the isolates with F126L/V125A appeared resistant to fluconazole (MIC ≥32 μg/mL). We also identified polymorphisms in S639 in the hotspot1 of the *FKS1* gene in 90% of the isolates with decreased susceptibility to micafungin. The most frequent mutation was S639P in 13 isolates from Clade IV (11 resistant to micafungin), and S639F and S639Y were found in micafungin-resistant isolates from Clade I and III (**Figure 4**).

Analysis of the distribution of *ERG11* copy number variation (CNV) revealed that of 304 isolates, 18 (6%) had either two or three copies. Of those 18 isolates, all were resistant to fluconazole and 17 (94%) were from Clade III (**Supplementary Figure 5**). Isolates within Clade III with 2 and 3 copies of *ERG11* had significantly higher MICs (*P* ≤ 0.05; Mann-Whitney test) to fluconazole than isolates with one copy (**Supplementary Figure 5**). Along with CNVs in *ERG11*, we found a total of six long regions (>40 kb) that showed increased copy number. Unlike CNVs in *ERG11*, these CNVs appeared in single isolates even in highly clonal clusters, with two isolates in each of Clades I, II, and IV (**Supplementary Figure 6**). While genes in these regions (between 23 and 125 genes in each region) have no direct relation with antifungal resistance, they might play a role in microevolution and *C. auris* adaptation to host stress. This includes genes associated with response to oxidative stress (*AOX2*, *HSP12*), iron assimilation (*FET33*, *FTR1*, *CHA1*, *FLC1*), cell wall and membrane integrity (*MNN2*, *ERG5*, *ERG24*), a transcription factor (*ZCF16*) and oligopeptide transporters associated with metabolic and morphologic adaptation and adherence (**Supplementary Table 2**). These data provide insight into the underlying molecular mechanisms of antifungal resistance and suggest that CNV could be a mechanism of strain variation in *C. auris*. Further exploration and monitoring of these traits are crucial to improve our understanding of *C. auris* diversity and control the expanding outbreak.

## Discussion

In this study, we used whole-genome sequencing to describe a global collection of *C. auris* isolates collected from patients and healthcare facilities between 2004 and 2018. We found that the four predominant clades are genetically distinct with strong geographic substructure in Clade IV. Using collection dates to estimate a molecular clock, we dated the origins of the four clades and confirmed the recent emergence of *C. auris*. Furthermore, we characterized mutations associated with antifungal resistance by clade, which varied between clades and country of isolation. While the clades appear largely clonal in species phylogenies and represent a single mating type, we found that they have distinct evolutionary histories and genome-wide patterns of variation. We provided a browsable version for *C. auris* genomic epidemiology through Microreact (28) to explore phylogeny, geographic distribution, timeline and drug resistance mutations (https://microreact.org/project/wfwXVjf9G).

In contrast to previous reports, we observed more phylogeographic mixing for *C. auris* (13). While we found that isolates from additional global regions can be clearly assigned to one of the four previously reported clades, we observed that three countries – Canada, Kenya, and United States – had isolates corresponding to multiple *C. auris* clades (**Figure 1B**). Additionally, isolates from multiple clades have been previously reported in Germany (19) and the United Kingdom (20). As travel has been previously shown to play a major role in the spread of *C. auris* (21), global travel of persons with prior healthcare exposures to *C. auris* has likely contributed to the observed phylogeographic mixing. Our analysis of the likely geographic origin of infections observed in new geographic regions is limited by incomplete travel history for most patients in this study set. We noted that the strongest geographic substructure was observed in Clade IV for isolates from Colombia, Panama and Venezuela, with additional distinct clades of isolates from Israel and United States (**Figure 1C**). This finding further supports evidence of rapid localized transmission in some of these countries (18, 21).

These results have confirmed prior findings from the analysis of a smaller data set where isolates in Clades I and IV had *MTL***a** and those in Clades II and III had *MTL*α (14). Although mating between *C. auris* clades has not been reported, it is concerning that the majority of countries reporting multiple *C. auris* clades have clades of opposite mating types. This is especially concerning in Kenya, where opposite mating types were observed in single healthcare facility experiencing ongoing transmission. In such a situation, it could be possible to have mixed infections of opposite mating types. If mating occurred, this would lead to increased genetic diversity and the possibility for enhanced virulence and exchange of drug resistance alleles. Continued efforts to characterize *C. auris* infections at the genomic level are essential for the rapid detection of potential *C. auris* hybrids.

Assessment of *C. auris* population structure by PCA and genome-wide F_ST_ analysis yielded no evidence for admixture between the major clades. The close relationship of Clades I and III is highlighted by the detection of regions with very low F_ST_ values, which suggest recent divergence or genetic exchange between these clades. Given that these regions were short and spread across the genome, we hypothesized that they are a result of incomplete lineage sorting rather than recent introgression events. We also observed variation in the average divergence between clades (D_XY_) along each chromosome. This may be due to genome rearrangements between the clades, whereby genomic areas exhibiting high D_XY_ levels, or low gene flow, arose in *C. auris* due to chromosomal rearrangements, which prevents recombination and supports high rates of genetic differentiation. Variation in chromosome number and size as measured by electrophoretic karyotyping as well as deletions, inversions and translocations detected by comparing genome assemblies of different *C. auris* clade isolates have been described in *C. auris* (14, 26, 27).

This global survey has provided a wider perspective of the mechanisms and the frequency of mutations associated with resistance to antifungal drugs. The presence of both resistant and susceptible isolates in the same populations along with the presence of genetically related isolates with different alleles of resistance genes indicate that the resistance in *C. auris* is not intrinsic and has been recently acquired. The most common mutation associated with azole resistance in Clades I and IV was *ERG11* Y132F; however, both clades also included genetically related isolates with *ERG11* K143R. In contrast, all fluconazole resistant isolates in Clade III carried *ERG11* F126L substitution. In addition to mutations in genes associated with drug resistance, we found that increase in copy number of *ERG11* is predominantly observed in Clade III again suggesting clade specific variation in mechanisms of azole resistance. All but three isolates with micafungin resistance had *FKS1* S639Y/P/F mutations. Taken together, these observations suggest recent emergence of antifungal resistance in *C. auris* populations, most likely in response to some unknown environmental change, such as increased use of azole antifungals in clinical practice, agriculture, or both.

By using a molecular clock, we estimated ages of the four clades by calculating time to the most recent common ancestor (TMRCA) of each clade. Our estimates demonstrated that Clade II was the oldest clade with TMRCA of 339 years, while Clade IV was the youngest with TMRCA of 34 years. Clade I and III isolates coalesce 140 and 175 years ago, respectively; however, in both Clade I or Clade III the clusters of isolates associated with ongoing drug resistant outbreaks worldwide, which display increased resistance to fluconazole and harbor mutations in *ERG11* associated with drug resistance, have emerged within the last 35 years. These results are consistent with the other population characteristics: even with the smallest sample set, Clade II had the highest genetic diversity compared to the three other clades and a positive TD indicative of the balancing selection, characteristics of an older population. Conversely, three other clades had low genetic diversity and negative TD consistent with the rapid emergence. Notably, the oldest *C. auris* isolate was collected from a patient in South Korea in 1996 (2) and no other strains were identified by searching the historic *Candida* culture collections. The absence of *C. auris* in culture collections prior to 1996 and a rapid emergence after 2012 suggest that this organism only recently emerged as a human pathogen and likely occupied a different ecological niche.

Other notable fungal outbreaks also have been estimated to be of recent origin. For example, the BdGPL lineage of the amphibian pathogen *Batrachochytrium dendrobatidis* was estimated to have arisen only ~100 years ago (29). While our reported mutation rate of 1.8695^e-5^ substitutions per site is consistent with that (5.7^e-5^; *R*^2^ = 0.37) reported in a previous study (17), the mutation rate over longer time spans than we sampled is likely slower. We used collection dates spanning from 2004 to 2018 to inform our estimate, and rates of molecular evolution measured over short time-scales tend to be overestimated, as some sites will be removed over time by natural selection (30). Therefore, the rate is more similar to a spontaneous mutation rather than an evolutionary substitution rate. If our mutation rate is substantially overestimated, the exact times of *C. auris* emergence and clade divergence would be older than we have estimated. We also acknowledge that utilizing only currently known isolates, which are highly similar within clades, provides a limited sampling of a larger source population, which may be also be undergoing sexual recombination. The identification and characterization of a wider population sample of *C. auris* will provide a higher resolution view of the nodes separating these major clades. However, as there is only speculation to date about potential associations or locations of such a source population, we suggest that the dates reported be used as a rough estimate that will need further evaluation when sources of additional diversity are identified.

Our molecular clock estimates demonstrate that nearly all outbreak-causing clusters from Clades I, III and IV originated 34-35 years ago in 1984-85. Such recent origin and nearly simultaneous detection of genetically distinct clades suggest that anthropogenic factors might have contributed to its emergence. Specifically: 1) In the 1980s, azole drugs first became widely used in clinical practice. The first azole topical antifungal drug, miconazole, was approved in 1971 followed by clotrimazole in 1972; both became widely used for treatment of superficial fungal infections in the late 1970s. In 1981, the first oral azole drug, ketoconazole, was released for treatment of systemic fungal infections (31). 2) In agriculture, the first azole fungicides, triadimefon and imazalil, were introduced in 1973, and by the early 1980s, ten different azole pesticide formulations were available. It has been demonstrated that azoles from the agricultural use can penetrate ground water and accumulate in soils. 3) Also noteworthy, our predicated emergence of *C. auris* as a human pathogen coincided with the early stages of AIDS epidemics; however, the wide use of antifungal drugs, such as fluconazole, for treatment of secondary fungal infections, had not started until the 1990s (32–34). Other anthropogenic factors might also have brought *C. auris* into contact with humans (35). Although the emergence of *C. auris* may be due to multiple factors, the coincidence between the introduction of azoles and *C. auris* emergence is intriguing and requires further investigation. Understanding processes that led to the emergence of *C. auris* in humans is important to prevent emergence of other drug resistant fungi and pathogens.

Although a recent study reported an isolate from a fifth clade isolated from a patient in Iran (16), all isolates in our collection were assigned to the previously described Clades I, II, III, and IV. This is noteworthy because isolates from neighboring Pakistan, Saudi Arabia, and United Arab Emirates were represented in the analysis. Indeed, this highlights the unique nature of the divergent Iranian *C. auris* case and advocates for increasing diagnostic capacity worldwide and continued phylogenetic studies to understand *C. auris* diversity.

While we have included a set of diverse isolates, they likely differ from a random sample of the *C. auris* population. The isolates were obtained by conventional sampling, and, therefore, our findings do not represent country-specific characteristics of *C. auris* molecular epidemiology. Wider sampling may also change the population structure and antifungal susceptibility profiles. Notably, at the time of this analysis, Clade V had not yet been discovered, highlighting the importance of further sampling and genomic characterization. Finally, since the environmental reservoir of *C. auris* remains unknown, our analysis is based solely on the analysis of clinical isolates; higher genetic diversity, deeper divergence times and different population structure is likely to occur in the natural populations of this fungus.

In conclusion, we have provided a comprehensive genomic description of a global *C. auris* survey representing 19 countries on six continents. Given that *C. auris* is a transmissible multidrug-resistant organism causing outbreaks of invasive infections in healthcare studies, an understanding of how *C. auris* is spreading, evolving, and acquiring resistance to antifungal drugs is essential for robust public health responses. Continued efforts to characterize the *C. auris* population, additional mechanisms of antifungal resistance, and environments conducive for mating between clades is critical.

## Methods

### Sample collection

We performed genomic analyses on sequences from 304 *C. auris* isolates. This collection included *C. auris* from 19 countries on six continents and were sampled from both *C. auris* cases and environmental surfaces from healthcare facilities where ongoing transmission was occurring (**Supplementary Table 1**). Samples from *C. auris* cases were derived from a variety of specimen source sites including sterile sites, such as blood, and non-invasive sites, such as respiratory tract or urine. All samples were a result of convenience sampling. For four countries (Colombia, Kenya, United States, and Venezuela) where more than 50 samples were available, 50 representative samples were selected by proportional random sampling: samples from each country were stratified by city, and then a subset was randomly selected proportionally from each strata.

### Sample preparation and whole-genome sequencing (WGS)

The sample collection comprised both publicly available sequences generated from previous studies and newly sequenced isolates (**Supplementary Table 1**). For newly sequenced isolates, except those from France, DNA was extracted using the ZR Fungal/Bacterial DNA MiniPrep kit (Zymo Research, Irvine, CA, USA). For isolates from France, DNA was extracted using NucleoMag plant kit extraction (Macherey-Nagel, Germany) in a KingFisherTM Flex system (Thermo Fisher Scientific). Genomic libraries were constructed and barcoded using the NEBNext Ultra DNA Library Prep kit for Illumina (New England Biolabs, Ipswich, MA, USA) and were sequenced on either the Illumina HiSeq 2500 platform (Illumina, San Diego, CA, USA) using the HiSeq Rapid SBS Kit v2 500-cycles or the MiSeq platform using the MiSeq Reagent Kit v2 500-cycles. For the two isolates from France, libraries were constructed using the Illumina Nextera Flex protocol and sequenced on an iSeq 100 to generate paired 150 bp reads.

### Variant identification

We used FastQC v0.11.5 and PRINSEQ v0.20.3 (36) to assess read quality and perform filtering for low quality sequences using “-trim_left 15 -trim_qual_left 20 -trim_qual_right 20 -min_len 100 -min_qual_mean 25 -derep 14”. Paired-end reads were aligned to the *C. auris* assembly strain B8441 (GenBank accession PEKT00000000.2; (14)) using BWA mem v0.7.12 (37). Variants were then identified using GATK v3.7 (38) using the haploid mode and GATK tools *RealignerTargetCreator*, *IndelRealigner*, *HaplotypeCaller* for both SNPs and indels, CombineGVCFs, GenotypeGVCFs, GatherVCFs, SelectVariants, and Variant Filtration. Sites were filtered with Variant Filtration using “QD < 2.0 ‖ FS > 60.0 ‖ MQ < 40.0”. Genotypes were filtered if the minimum genotype quality < 50, percent alternate allele < 0.8, or depth < 10 (https://github.com/broadinstitute/broad-fungalgroup/blob/master/scripts/SNPs/filterGatkGenotypes.py). Genomic variants were annotated and the functional effect was predicted using SnpEff v4.3T (39). The annotated VCF file was used to determine the genotype of known mutation sites in *ERG11* and *FKS1*. To determine the mating type locus (*MTL***a** and *MTL*α) in each isolate, the average read depth at the locus was computed from the aligned bam file and normalized by the total coverage depth.

### Phylogenetic and phylodynamic analysis

For phylogenetic analysis, sites with an unambiguous SNP in at least 10% of the isolates (*n* = 222,619) were concatenated. Maximum likelihood phylogenies were constructed using RAxML v8.2.4 (40) using the GTRCAT nucleotide substitution model and bootstrap analysis based on 1,000 replicates. Phylogenetic analysis was also performed for each clade using subsets of the entire VCF and visualized using iTOL (41).

For phylodynamic analysis, we assessed for a temporal signal using a set of isolates from Kenya Clade III using TempEst v1.5.3 (42) to quantify and to estimate an initial mutation rate from the R-squared value. Bayesian phylogenies were generated using BEAST v1.8.4 (43) under strict molecular clock (both lognormal and exponential priors). In addition, we applied both Bayesian Skyline coalescent and Coalescent Exponential, and a GTR nucleotide substitution model. We obtained similar results using the molecular rate estimated for a *C. auris* outbreak un United Kingdom (17). Specimen collection dates (month and year) were used as sampling dates; the month of June (year midpoint) was assigned for samples where month was unknown. Bayesian Markov chain Monte Carlo (MCMC) analyses were run for 20 million steps using an unweighted pair-group method with arithmetic mean (UPGMA) tree as a starting tree and MCMC convergence was explored using Tracer v.1.7.1 (44). We generated a maximum clade credibility tree with TreeAnnotator v1.8.4 after discarding 47% as burn-in, and we visualized phylogenies using FigTree v1.4.4.

### Population genomic analyses

Population genomic analysis was performed using gdsfmt v1.14.1 (34), SNPRelate v1.12.2 (45), and the PopGenome v2.6.1 (46) R packages. Genome-wide nucleotide diversity (π), Tajima’s D (TD), fixation index (F_ST_) and pairwise nucleotide diversity (D_XY_) were calculated and plotted per scaffold in 5 kb sliding windows. Genome-wide calculations are the average of all 5 kb windows for each metric. Genomic regions that exhibit copy number variation (CNV; deletions and duplications) were identified using CNVnator v0.3 (47) (genomic windows > 1 kb showing significant variation *P*-value < 0.01).

### Antifungal susceptibility testing (AFST)

AFST was performed on 294/304 (97%) isolates. The majority (N=270, 90%) of isolates were tested at the U.S. Centers for Disease Control and Prevention (CDC) as outlined by Clinical and Laboratory Standards Institute guidelines. Custom prepared microdilution plates (Trek Diagnostics, Oakwood Village, OH, USA) were used for fluconazole and the echinocandin micafungin. Interpretive breakpoints for *C. auris* were defined based on a combination of those breakpoints which have been established for other closely related *Candida* species, epidemiologic cutoff values, and the biphasic distribution of minimum inhibitory concentrations (MICs) between the isolates with and without known mutations for antifungal resistance. Resistance to fluconazole was set at ≥32 μg/mL and at ≥4 μg/mL for micafungin. Amphotericin B was assessed by Etests (BioMerieux), and resistance was set at ≥2 μg/mL. For isolates not tested at the CDC, similar methods were employed and described previously (17, 19, 48). As there are no currently approved breakpoints for *C. auris*, for this manuscript breakpoints were set at ≥32 μg/mL for fluconazole, >1 μg/mL for amphotericin B, and ≥4 for micafungin. These MIC values were based on a combination of the wild type distribution (those isolates with no mutations) and PK/PD analysis in a mouse model of infection (49).

## Data and resource availability

All Illumina sequence generated by this project is available in the NCBI SRA under BioProjects PRJNA328792, PRJNA470683, PRJNA493622, and PRJNA595978. The phylogenetic tree has been deposited in Microreact (https://microreact.org/project/wfwXVjf9G). A set of isolates are available from the CDC and FDA Antimicrobial Resistance (AR) Isolate Bank (https://www.cdc.gov/drugresistance/resistance-bank/index.html).

### Ethics

This project was reviewed by CDC IRB as part of the broader human subjects protocol for the Mycotic Diseases Branch, CDC.

## Acknowledgements

The authors thank Anuradha Chowdhary, Joveria Farooqi, Nelesh Govender, Mathew Fisher, and Koichi Makimura for their continued support and sharing isolates, and Mathew Fisher and Angela Early for comments on the manuscript. We also thank Jacques Meis for coordinating isolate acquisition; Javier Peman and Olga Rivero-Menendez for support with the Spanish isolates; Erika Santiago, Jovanna Borace and Angel Cedeño from Hospital Santo Tomas; and Soraya Salcedo, Adriana Marín, Carmen Varón, Nohora Villalobos, Jairo Perez, Julian Escobar for participating in the collection of strains and data from Colombian institutions. This project has been funded in part with Federal funds from the National Institute of Allergy and Infectious Diseases, National Institutes of Health, Department of Health and Human Services, under award U19AI110818 to the Broad Institute. CAC is a CIFAR fellow in the Fungal Kingdom Program. This work was also made possible through support from the Advanced Molecular Detection (AMD) initiative at CDC.

## Disclaimer

The use of product names in this manuscript does not imply their endorsement by the U.S. Department of Health and Human Services. The finding and conclusions in this article are those of the authors and do not necessarily represent the views of the Centers for Disease Control and Prevention.

