## Supplementary Figures for "Tracing the evolutionary history and global expansion of *Candida auris* using population genomic analyses"

[1]Mycotic Diseases Branch, Centers for Disease Control and Prevention, U.S. Department of Health and Human Services, Atlanta, GA, 30329 USA; [2]Broad Institute of MIT and Harvard, Cambridge, MA, 02142 USA; [3] Department of Pathology, Aga Khan University Hospital, Nairobi, Kenya , [4]Institut Pasteur, Molecular Mycology Unit, CNRS UMR2000, National Reference Center for Invasive Mycoses and Antifungals (NRCMA), Paris, France; [5]Laboratoire de Parasitologie-Mycologie, Hôpital Saint-Louis, Groupe Hospitalier Lariboisière, Saint-Louis, Fernand Widal, Assistance Publique-Hôpitaux de Paris (AP-HP); [6]Université Paris Diderot, Université de Paris, Paris, France ; [7]Mycology Reference Laboratory, National Centre for Microbiology, Instituto de Salud Carlos III, Carretera de Majadahonda a Pozuelo Km. 2 28220, Majadahonda, Madrid, Spain; [8]Department of Pathology and Laboratory Medicine, King Faisal Specialist Hospital and Research Centre, Riyadh, Saudi Arabia; [9]Hospital Santo Tomás, Panama City, Panama; [10]Infectious Diseases Unit, Tel Aviv Sourasky Medical Center, and the Sackler Faculty of Medicine, Tel Aviv University, Tel Aviv, Israel; [11]National Microbiology Laboratory, Public Health Agency of Canada, Winnipeg, MB, R3E 3R2, Canada; [12]Department of Infectious Diseases, School of Medicine, Universidad del Zulia, Maracaibo, Venezuela; [13]Institut Pasteur, Molecular Mycology Unit, CNRS UMR2000, Paris, France; [14]Grupo de Microbiología, Instituto Nacional de Salud, Calle 26 # 51-20, Bogotá, Colombia; [15]Department of Microbiology, PathWest Laboratory medicine FSH Network, Fiona Stanley Hospital, Murdoch, Australia; [16]Department of Microbiology, PathWest Laboratory Medicine FSH Network, Fiona Stanley

Hospital, Murdoch; [17]Department of Infectious Diseases, Fiona Stanley Hospital, Department of Microbiology; [18]Infectious Diseases, Royal Perth Hospital, Perth, WA, Australia; [19]Faculty of Health & Medical Sciences, University of Western Australia, Crawley, WA, Australia; [20]German National Reference Center for Invasive Fungal Infections NRZMyk, Leibniz Institute for Natural Product Research and Infection Biology – Hans-Knöll-Institute, Jena, Germany; [21]University of Würzburg, Institute for Hygiene and Microbiology, Würzburg, Germany

Current addresses; XL: Advanced Institute of Information Technology, Peking University, Hangzhou, ZJ, China. 311215.

Running Head: Genomic analysis of *Candida auris* worldwide

*N.A.C. and J.F.M. contributed equally to this work. Co-first authorship order was assigned alphabetically.

[#]A.P.L. and C.A.C. contributed equally to this work. Co-senior authorship was assigned based on the co-first authorship and the equal contributions of both groups.

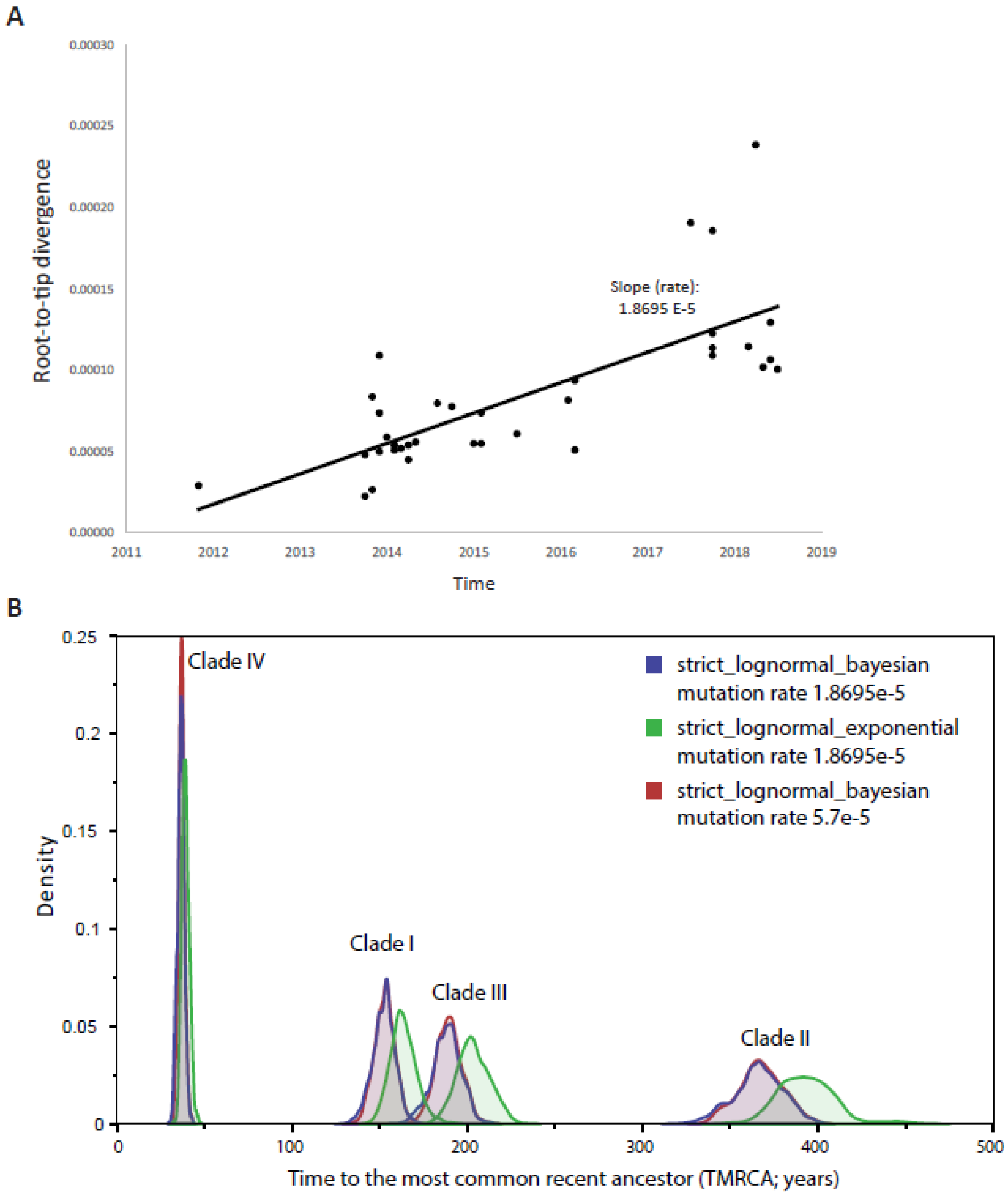


**Supplementary Figure 1. (a)** Root-to-tip regression analysis performed using a maximum likelihood tree of *C. auris* genomes from Kenyan isolates clustering to Clade III. R-squared value = 0.55. **(b)** Marginal posterior distributions for the date of the most recent common ancestor (TMRCA) of *Candida auris* Clades I, II, III and IV, analysis performed by BEAST under distinct models and mutation rates.

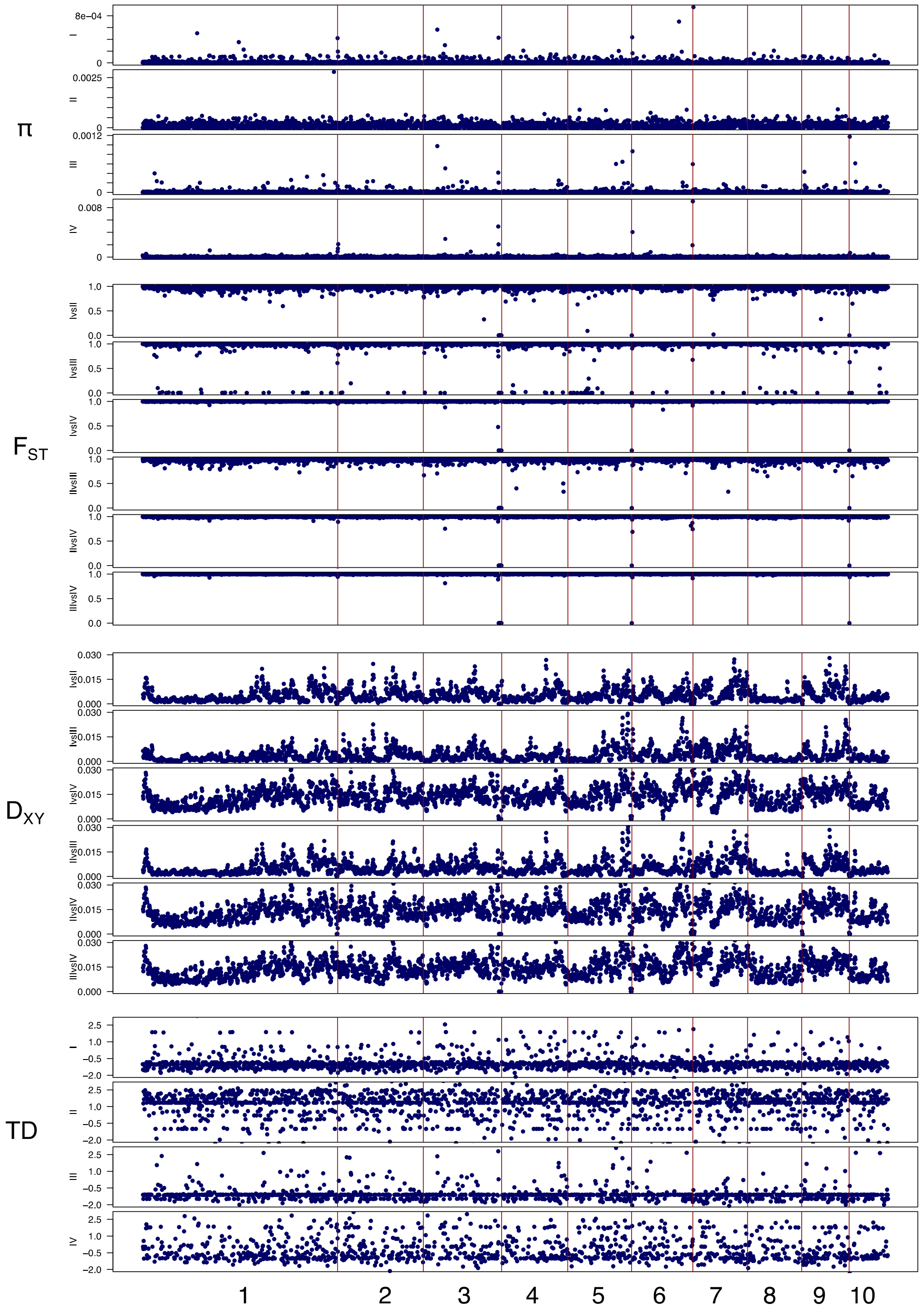


**Supplementary Figure 2. Population genomic analysis of *C. auris*.** Genome-wide nucleotide diversity (π), Tajima's D (TD), fixation index ($F_{ST}$) and pairwise nucleotide diversity ($D_{XY}$) were calculated per scaffold in 5 kb sliding-window and plotted across the genome. Plots depict the ten largest scaffolds of the B8441 reference genome.

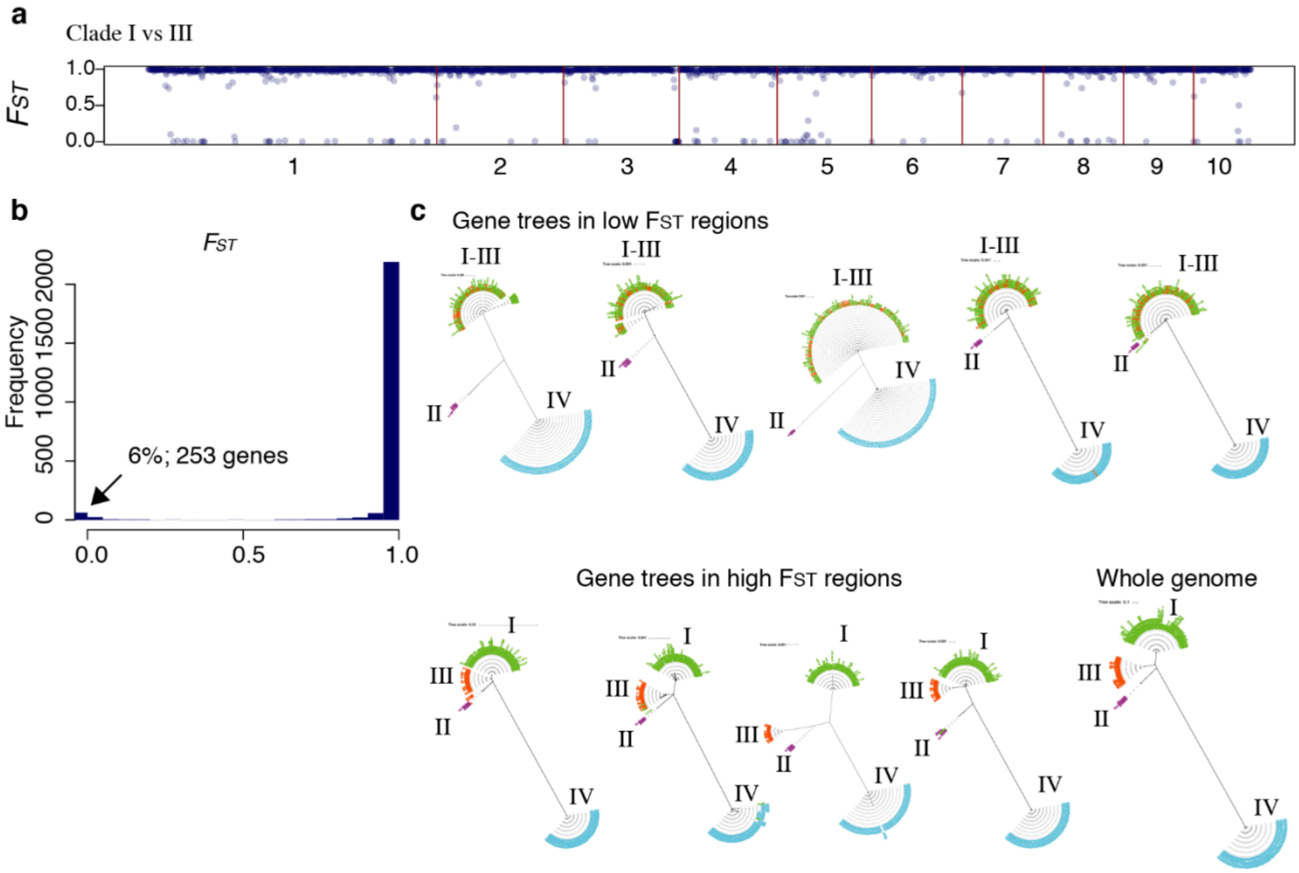


**Supplementary Figure 3. Genome-wide FST analysis of the two most closely related clades (Clades I and III). (a, b)** Genome-wide FST analyses revealed substantial interspecific divergence and reproductive isolation between *C. auris* Clades I and III and small regions with FST values close to zero. **(c)** Phylogenetic analysis of selected genes revealed that these regions in isolates from Clades I and III are intermixed in a monophyletic clade.

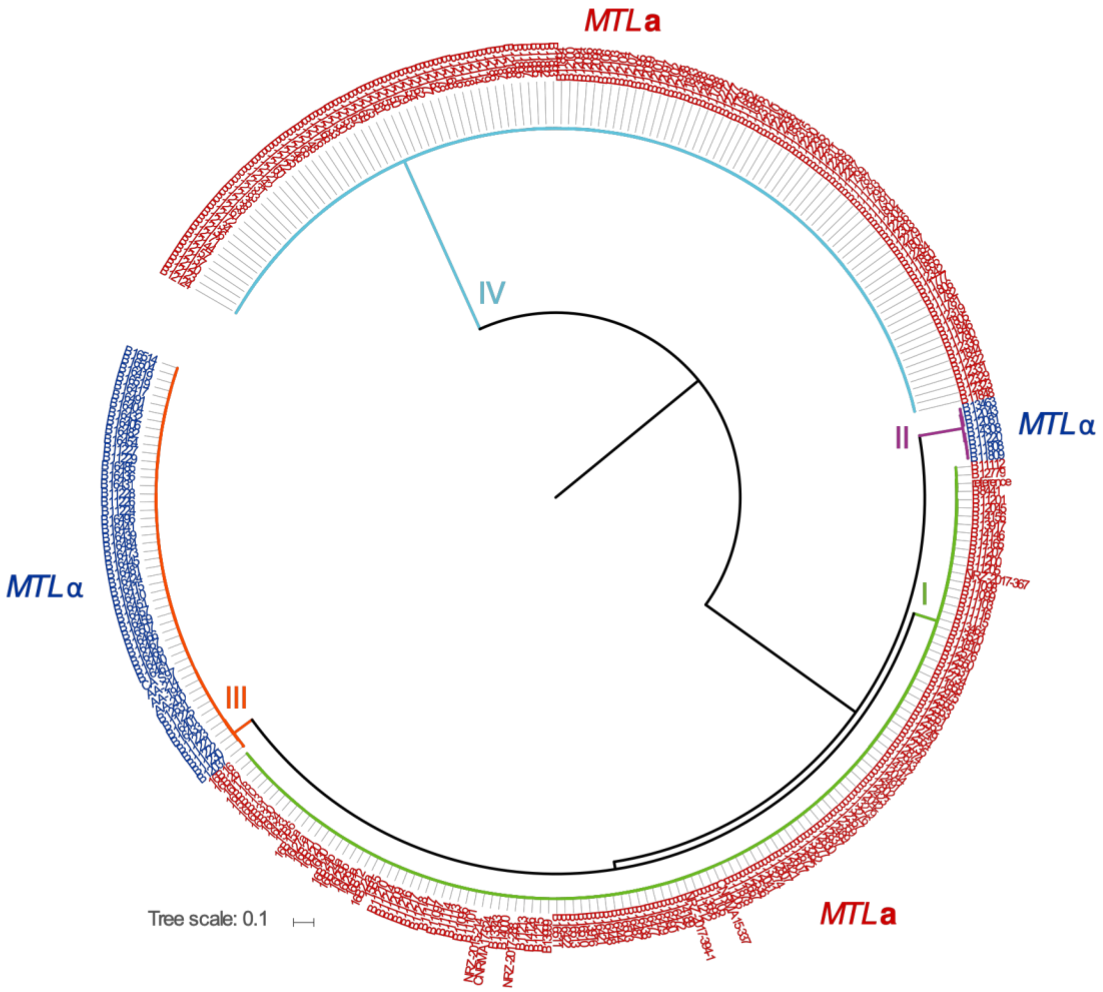


**Supplementary Figure 4. Mating-type locus classification in *C. auris*.** Phylogenetic tree of 304 *C. auris* isolates. Branches are colored by clade and isolate labels are colored by mating-type locus (*MTL***a**: red and *MTL*α: blue).

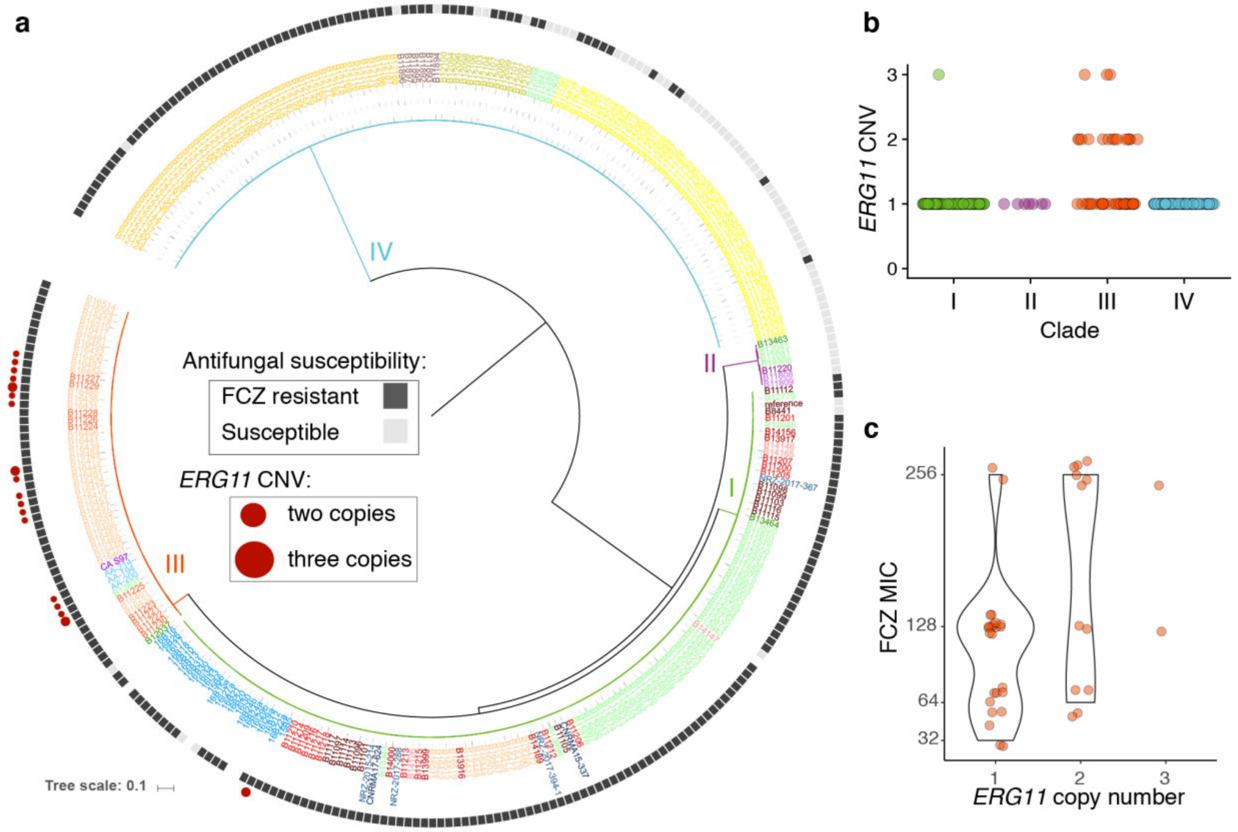


**Supplementary Figure 5. Copy number variation (CNV) of *ERG11* in *Candida auris*. (a)** Phylogenetic tree detailing clade, country of origin, susceptibility to fluconazole (FCZ resistant or susceptible). Isolates that have more than one copy of lanosterol 14-alpha-demethylase (*ERG11*) are denoted with a red circle. **(b)** Plot shows the frequency of isolates that have 1, 2 or 3 copies or *ERG11* by clade. **(c)** Plot depicts fluconazole minimal inhibitory concentration (MIC) results for 38 isolates from Clade III (35 from Kenya and 3 from Spain) grouped into copy number of *ERG11*.

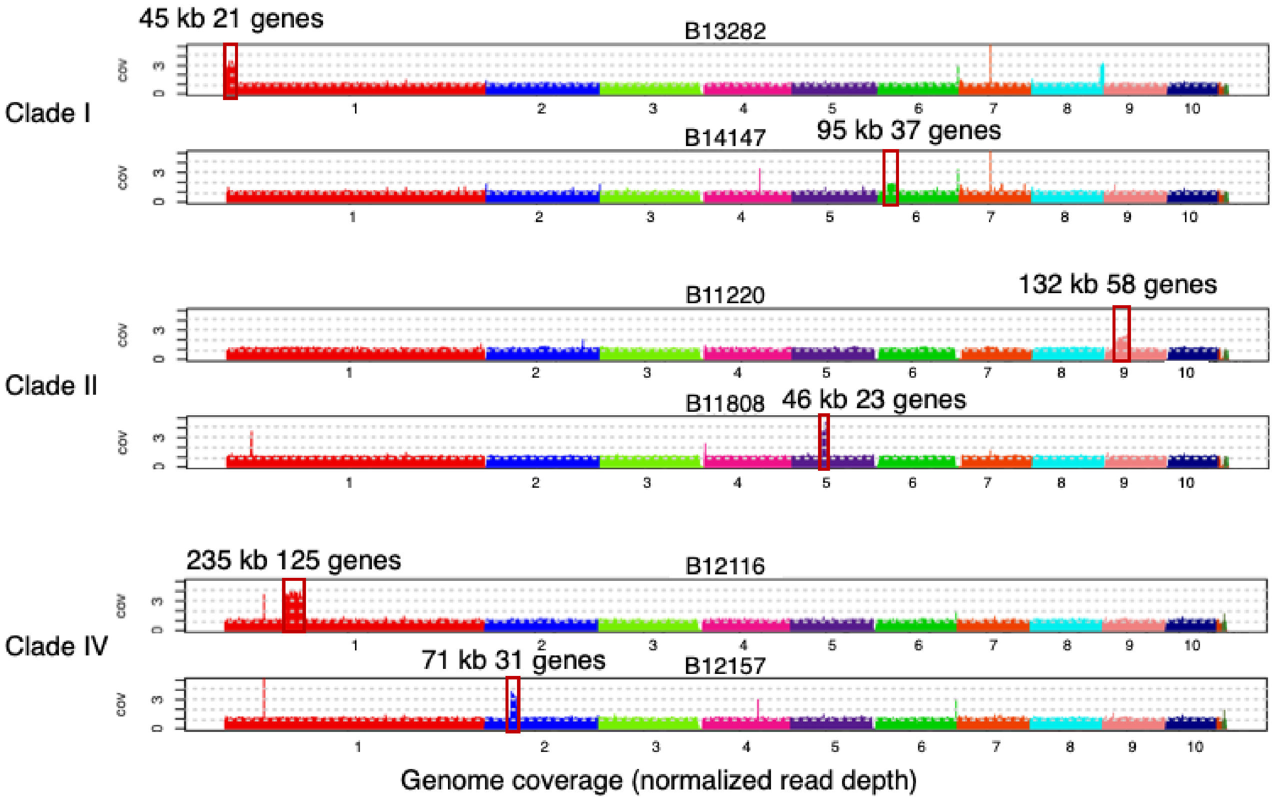


**Supplementary Figure 6.** Other regions with copy number variation (CNV) in *C. auris* population.
